# *Plankton Planet*: ‘seatizen’ oceanography to assess open ocean life at the planetary scale

**DOI:** 10.1101/2020.08.31.263442

**Authors:** Colomban de Vargas, Thibaut Pollina, Sarah Romac, Noan Le Bescot, Nicolas Henry, Calixte Berger, Sébastien Colin, Nils Haëntjens, Margaux Carmichael, David Le Guen, Johan Decelle, Frédéric Mahé, Emmanuel Malpot, Carole Beaumont, Michel Hardy, *the planktonauts*, the *Plankton Planet team*, Damien Guiffant, Ian Probert, David F. Gruber, Andy Allen, Gabriel Gorsky, Mick Follows, Barry B. Cael, Xavier Pochon, Romain Troublé, Fabien Lombard, Emmanuel Boss, Manu Prakash

## Abstract

In every liter of seawater there are between 10 and 100 billion life forms, mostly invisible, called plankton, which form the largest and most dynamic ecosystem on our planet, at the heart of global ecological and economic processes. While physical and chemical parameters of planktonic ecosystems are fairly well measured and modelled at the planetary scale, but biological data are still scarce due to the extreme cost and relative inflexibility of the classical vessels and instruments used to explore marine biodiversity. Here we introduce ‘*Plankton Planet*’, an initiative whose goal is to merge the creativity of researchers, makers, and mariners to (*i*) develop frugal scientific instrumentation and protocols to assess the genetic and morphological diversity of plankton life, and (*ii*) organize their systematic deployment through fleets of volunteer sailors, fishermen, or cargo-ships to generate comparable and open-access plankton data across global and long-term spatio-temporal scales. As proof-of-concept, we show how 20 crews of sailors (“planktonauts”) were abl to sample plankton biomass from the world surface ocean in a single year, generating the first citizen-based, planetary dataset of plankton biodiversity based on DNA barcodes. The quality of this dataset is comparable to that generated by *Tara* Oceans and is not biased by the multiplication of samplers. This dataset has unveiled significant genetic novelty and can be used to explore the taxonomic and ecological diversity of plankton at both regional and global scales. This pilot project paves the way for construction of a miniaturized, modular, evolvable, affordable and open-source citizen field-platform that will allow systematic assessment of the eco/morpho/genetic variation of aquatic ecosystems across the dimensions of the Earth system.

## Global surveys of marine plankton life: urgency and challenges

The ocean contains 97% of all water on our planet. In every liter of seawater there are between 10 and 100 billion, mostly invisible planktonic life forms. These form a continuous global ecosystem that generates approximately half of planetary oxygen, sustains the large majority of marine life, and regulates atmospheric CO_2_ and climate. Understanding and modelling the structure, dynamics, and evolution of global plankton populations is critical for predicting the future of our biosphere and learning how to live in symbiosis with our spaceship, the Earth.

Plankton populations comprise, like in terrestrial biomes, organisms from across the tree of life (viruses, bacteria, archaea, protists, and animals) which interact in complex networks (*1, 2*) that are also shaped by ocean currents and associated physico-chemical environmental parameters (*3, 4*) – i.e. the *seascape* (*5*). But in contrast to terrestrial ecosystems, there are no plants and trees in the plankton - primary production is driven by a large and ancient diversity of photosynthetic bacteria and protists (called phytoplankton) - and the pelagic ecosystem is much more dynamic in terms of both organism life cycle and strategies (e.g. mixotrophy) (*6*) and transport (advection and mixing). The self-organization of local plankton biota into complex ecosystems (*7*) determines their impact on the carbon cycle. For instance, in some regions and seasons, blooms of relatively large cells with mineral components lead to a vigorous sinking flux of organic carbon into the deep sea (*8, 9*). Overall, these fundamental properties of the plankton ecosystem make it arguably the most reactive and proactive compartment of the biosphere to climate change and pollution. Changes in plankton communities can have dramatic effects on global biogeochemical cycles (e.g. *10*), climate (e.g. *11, 12*), and major human societal and economic activities (i.e. fisheries, aquaculture, tourism, etc. (*13, 14*).

Current models aimed at predicting global ocean ecological changes (e.g. (*15, 16*)) are fairly well constrained in terms of physics and chemistry, but are by comparison heavily oversimplified and unrealistic in terms of biology. In fact, they simply lack good quality, high resolution data on the nature and dynamics of oceanic plankton biodiversity *at a planetary scale*. While quantitative global data on ocean physics and biogeochemistry are abundantly available by satellites (e.g. *17*), *in situ* floats (e.g.*18, 19*)) and sail drones (e.g. *20*), as well as research (e.g. *21*) and citizen (e.g. *22*) vessels, standardized biological data are still scarce due to the significant challenge of sampling and assessing complex communities of fragile plankton in a harmonized and comparable manner (*23*). Despite the availability of a century’s worth of recorded data on oceanic plankton ((*24, 25*), and see for instance **Error! Hyperlink reference not valid**. some areas of the ocean (including much of the south and tropical Pacific Ocean) are almost devoid of any biological observations and we do not yet have an informed vision of the global distribution and variation of plankton communities.

The largest and longest homogenous survey of plankton life has been conducted using the Continuous Plankton Recorder (CPR), a visionary plankton-scroll instrument created by Sir Alister Hardy in 1931 (*26*), and towed since then behind ferries and cargo ships, particularly in the North Atlantic, North Pacific, and Southern Ocean south of Australia (*27*). The CPR database is currently the only basin-wide standardized historical record of ocean plankton life, and it has given rise to keystone studies describing and modelling basin-scale dynamics of plankton community over time and global climate change (e.g. *13, 28, 29*). However, CPR data also has major drawbacks: (*i*) the instrument mainly recovers zooplankton >300µm, thus missing most plankton diversity (see below), (*ii*) it is rather destructive for soft or gelatinous taxa, (*iii*) data analyses rely on taxonomic experts identifying and counting a restricted number of morpho-taxa, (*iv*) the formalin preservation of the silk severely limits subsequent light microscopy observations for some groups and analysis of nucleic acids in general, and (*v*) data come from ships navigating over a restricted number of (mostly northern) commercial routes. CPR data are therefore semi-quantitative and taxonomically and geographically limited.

Over the past 20 years, the revolution in environmental DNA/RNA sequencing has stimulated a new era of global-scale open ocean cruises led by molecular and cellular biologists, notably the ‘Global Ocean Sampling’ (GOS - *30*), *Tara* Oceans (*31*), and Malaspina (*32*) expeditions. Interestingly, the first two expeditions were private or semi-private enterprises undertaken by sailing boats. The *Tara* Oceans sailing expedition was the longest (2009 – 2013) and most comprehensive: its team developed an eco-systems biology strategy to explore plankton diversity from genes to communities, from viruses to animals, and across coarse but planetary spatial and seasonal scales (*33*). The combination of *standardized* DNA metabarcoding (*34, 35*), metagenomic (*36, 37*) and metatranscriptomic (*38, 39*) datasets has unveiled the basic structure of plankton taxonomic and metabolic diversity (*33, 40*), generating hypotheses about its interactions (*1*), dynamics in the seascape (*3*), and role in emerging ecosystem functions such as the carbon pump (*2, 41*). Notably, the *Tara* Oceans team discovered that the great majority of plankton biodiversity is found in organismal size fractions <100µm, and above all in eukaryotes rather than viruses or prokaryotes (*34, 38*).

Given the enormous local (e.g. plastics, pollutants (*42, 43*)) and global (e.g. deoxygenation, warming, acidification, freshening, ocean circulation changes (*44, 45*)) anthropogenic pressures on our ocean, we urgently need comprehensive global plankton surveys that merge the spatio-temporal sampling power of the CPR with the systems-biology approach of *Tara* Oceans. Only application of a comprehensive and homogenous measure of plankton life from micro-to meso-to global ocean scales (*7, 23, 46*) will provide the data necessary not only to unveil fundamental principles of ecology and evolution of marine life at the ecosystem level of organization, but also to feed mathematical models of the ocean system that integrate physical, chemical, and biological processes.

## Plankton Planet: toward a participative Oceanography 3.0

However, this endeavor is hindered by the extreme cost and limited logistical flexibility of classical oceanographic research vessels and instruments, together with the current impossibility to use autonomous samplers (e.g. floats) to sample the biocomplexity of plankton and generate high-quality data from it. In this context, the thousands of citizen sailing boats (*47*), professional sailing yachts, the >50,000 cargo ships (https://www.ics-shipping.org/shipping-facts/shipping-and-world-trade) and about the same number of global fishing vessels (https://globalfishingwatch.org/datasets-and-code/vessel-identity/) which are navigating the world ocean every day represent an outstanding opportunity.

With the miniaturization of sequencing (e.g. *48*) and imaging (e.g. *49*) devices, the power of cloud computing and artificial intelligence, and the possibilities offered by a participative science engaging actors and volunteers across disciplines and societal/economic activities (*22, 50*), five years ago we - a team of researchers, engineers, makers and sailors from France, the US, and New Zealand - started to develop a frugal approach and effective protocols to sample the world ocean plankton for the production of high quality eco/morpho/genetic data in collaboration with recreational and professional mariners. We created ‘*Plankton Planet*’ (P2, https://planktonplanet.org), an international initiative that promotes long-term citizen-based sampling of marine plankton to assess the biodiversity and health of the world’s open and coastal oceans (see P2 vision & mission, box 1).

### BOX 1.

**Plankton Planet Vision & Mission**

**Vision:** to harness the creativity of mariners, scientists, and makers, for global evaluation of ocean health, biodiversity, and evolution.

**Mission:** to develop user-friendly affordable tools to underpin a sustainable citizen Oceanography 3.0 providing critical new knowledge on global plankton morphology, genetics, and ecology, that will be universally accessible and feed mathematical modelling toward a predictable ocean.

The Oceanography 3.0 is independent and universal, based on interconnected individuals and instruments.

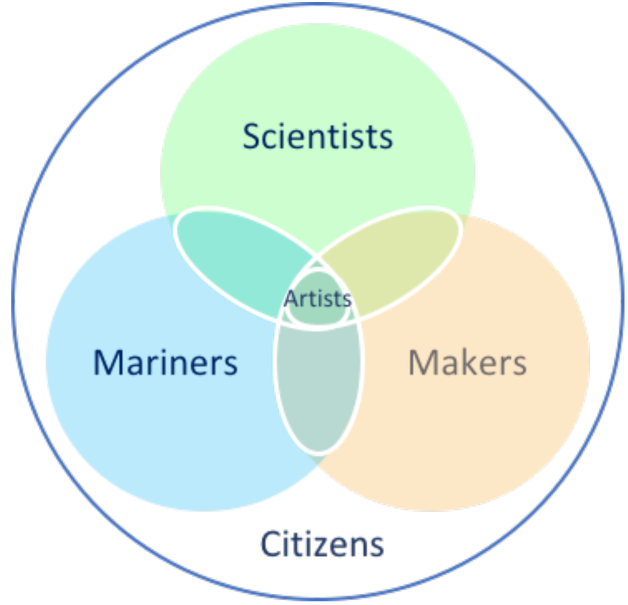

We strongly believe that mariners, makers, and researchers share a passionate curiosity and will to explore and discover their environment, such that erasing the boundaries between these worlds will generate the necessary synergies to achieve our objectives. Our practical goal is to co-construct a new generation of frugal yet robust scientific instruments and protocols, which will allow all volunteer mariners – from recreative or professional sailors to fishermen or cargo-ship crews – to collect comparable eco-morpho-genetic data on plankton diversity and abundance at a planetary scale, a concept coined

In this paper, we present the first steps of this cost-effective, eco-friendly, and society-engaging Oceanography 3.0. We show how the recruitment of the first 20 crews of citizen sailors – the planktonauts-, equipped with a simple kit to sample plankton for DNA-metabarcoding, were able to help generate a scientifically sound, planetary dataset of plankton biodiversity in less than a year. We finally discuss recent developments (see also (*51*)), as well as the ‘Plankton Planet’ perspectives toward the large-scale deployment of integrated, affordable and portable ‘field-eco-scopes’ for long-term assessment of ocean life and ecosystems at an unprecedented level of sensitivity.

## Plankton Planet: Proof-of-Concept

Between November 2014 and January 2016, with a total budget of $70,000, we provided proof-of-concept for the P2 vision and mission (Box 1). Our primary goal was to demonstrate that, with the goodwill of mariners, we can sample ocean waters from the entire planet and obtain high quality plankton DNA data for global ecological analyses. We designed a general functional strategy (Sup. Fig.1) based on robust methods linking affordable and user-friendly instruments for on-board citizen plankton sampling to cutting-edge DNA sequencing and bioinformatic pipelines developed previously in *Tara* Oceans (*34*).

### Frugal and global plankton sampling by citizen-sailors (‘planktonauts’)

#### The P2 PlanktoKit

We first assembled a simple plankton sampling kit including a small net to collect plankton (>20μm) and a manual pumping system to rapidly transfer the freshly collected plankton onto a filter membrane (Sup. Fig. 2). Of note, this sampling protocol does not rely on toxic chemicals or electricity, which are typically required in a regular laboratory. In order to avoid the need for high-energy storage of the filter membranes (plankton samples) in freezers on board, as well as complex frozen-shipping to the lab, the planktonauts were asked to place the filter on a clean aluminum cove and carefully heat-dry the filter membranes in a pan on the boat gas-cooker, store the dried plankton samples in labeled zip-lock bags with granular desiccants, and send them to the lab via standard postal services (Sup. Fig. 3, Sup. Text 1).

**Figure 1.**
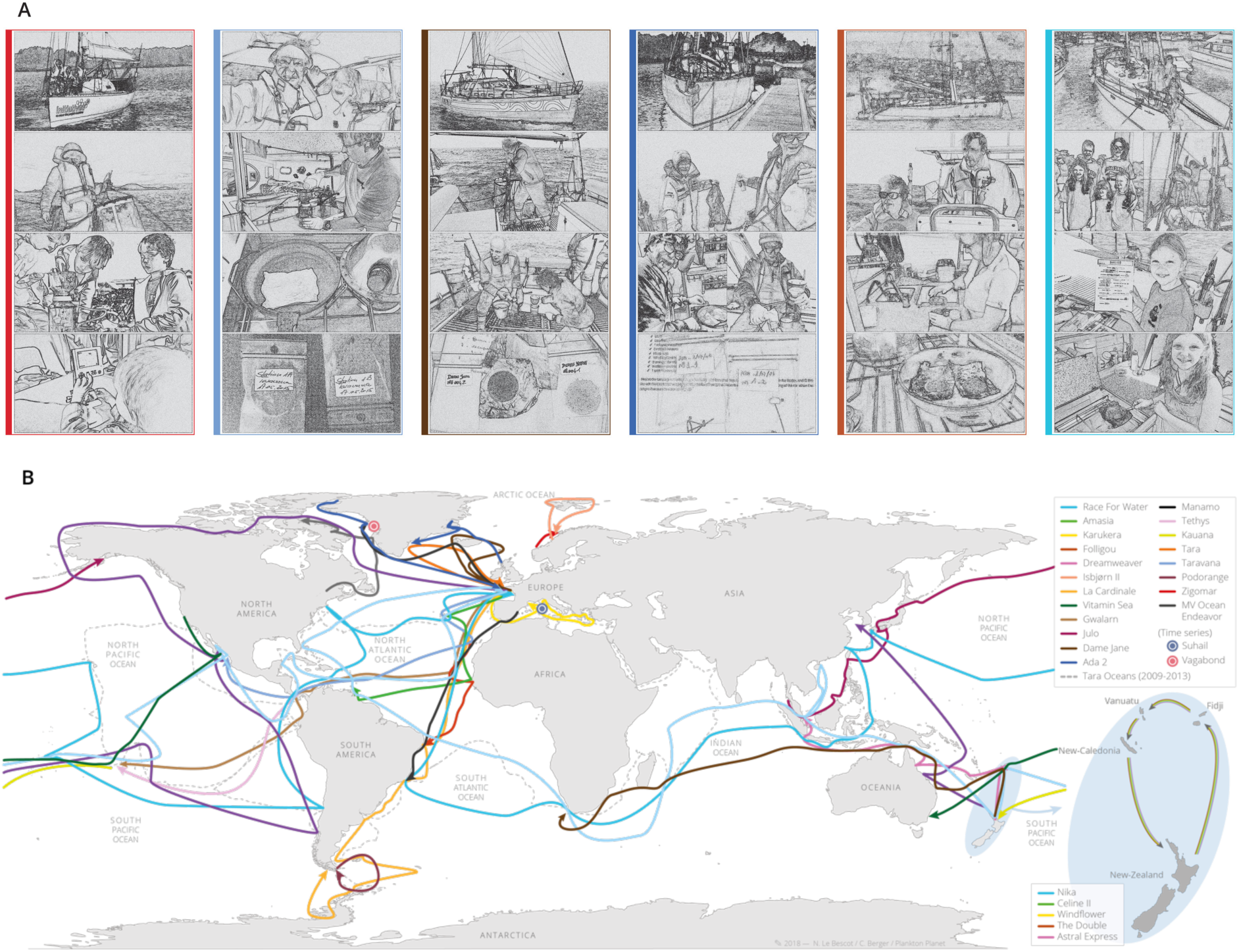
Planktonauts and their routes across the world ocean. **A**. Examples of pictures (sketch-like filtered) sent by the planktonauts illustrating their actions at sea: towing and recovering the P2 plankton net, filtering, drying, and storing plankton samples, recording contextual data and observing plankton through a microscope. **B**. Routes of the main 20 (of 27) pioneer planktonaut crews recruited during the pilot phase of the project, selected to maximize the geographic coverage and oceanographic and sampling conditions. Note the 5 boats from New Zealand who participated to a rally (May to November 2015) organized in collaboration with the ‘Island Cruising Association’ (http://www.islandcruising.co.nz).

**Figure 2.**
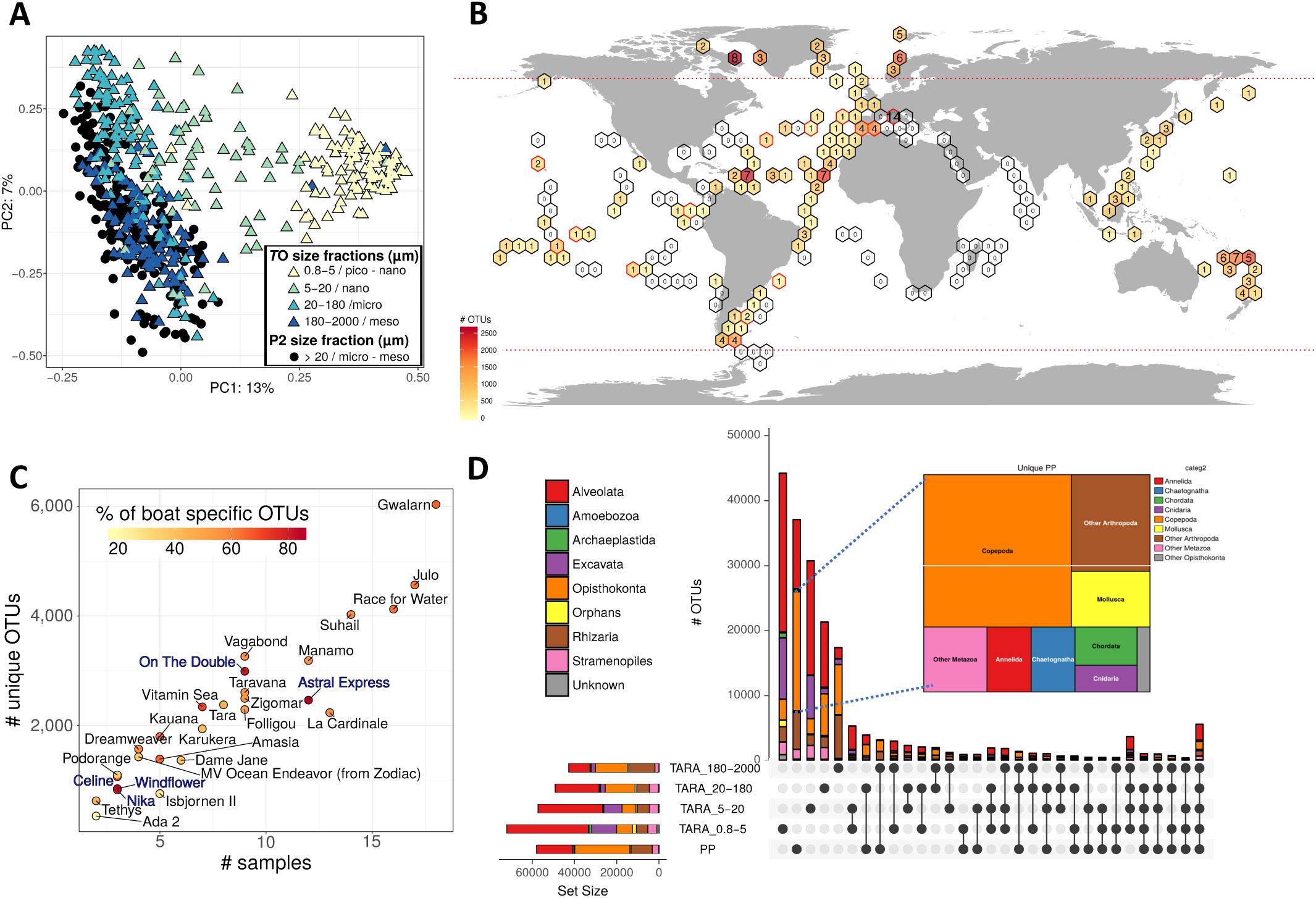
The extent, novelty, and community composition of P2 metabarcoding data. **A**. Grouping of P2 (black dots) and *T*O (colored triangles) plankton communities from surface water according to taxonomic compositional similarity (PCA of Hellinger standardized abundances). Colors correspond to the different plankton size-fractions sampled in *T*O. **B**. Geographic distribution and novelty of P2 sequenced samples as compared to *T*O samples. Sampled sites are aggregated in pre-defined geographic area (hexagons) for readability. Hexagons containing P2 samples are filled with a color gradient corresponding to the number of rDNA OTUs that were not detected in TO (see colored scale). Number inside the colored hexagons indicate the number of samples collected in the area. Empty hexagons are area with only *T*O samples. Hexagons with red-line borders are area with both P2 and *T*O samples. Horizontal dotted red lines indicate the Northern and Southern 60 Degree latitudes; only samples comprised between these lines were kept for ecological comparative analyses between P2 and TO data. **C**. Novel plankton diversity (rDNA OTU) uncovered by each boat. Each dot represents a boat, with its position along the X and Y axes corresponding, respectively, to the number of samples collected, and the number of OTUs recovered by the boat that were not detected in TO. The color-gradient indicates the % of novel P2 OTUs that are unique to the particular boat. Note the higher values for the New Zealander crews (blue characters). **D**. UpSetR plot displaying the taxonomic richness, divided by eukaryotic super-groups, shared between P2 (>20μm) and *T*O size-fractionated samples. The horizontal bars show each individual complete dataset, while vertical bars correspond to the number of OTUs shared between particular datasets (intersections given by the dots under the vertical bars). The tree map shows the taxonomic composition of the 18,430 unique Opisthokonta OTU unveiled in P2, with a dominance (46%) of copepods.

**Figure 3:**
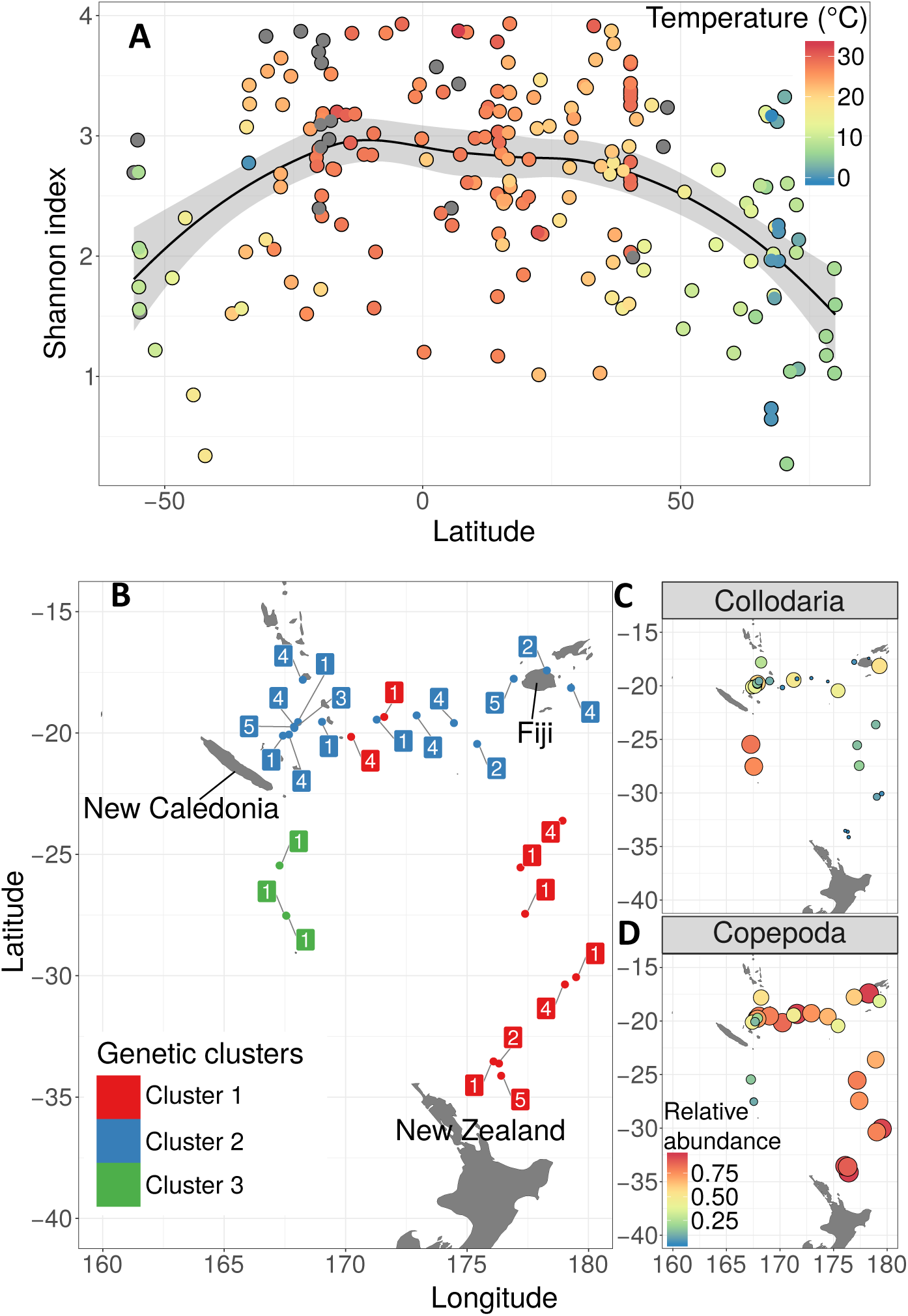
P2 insights into plankton ecology. **A**: *Global ocean latitudinal diversity gradient.* Scatter plot showing the evolution of the Shannon index (alpha diversity) of all P2 samples (colored dots) along latitudes. The color gradient is related to the sea surface temperature measured by the planktonauts during plankton sampling. LOESS regression curve^42^ fitting the data. **B-D**: *Plankton biogeography at the scale of a navigation loop* (the ‘kiwi loop’, around New Zealand, Fiji and New Caledonia, Fig. 2B). **B:**The samples from this loop (colored dots) were clustered into three groups based on their OTU composition (Jaccard distance) using the Partitioning Around Medoids (PAM) algorithm ^43^. See Sup. Fig. 9 for details. Two samples from the *Nika* boat, considered as outliers (and collected very close to the coast), were excluded from the analysis. Squared numbers close to each sample site correspond to the sampling boat (1: Astral Express; 2: Celine; 3: Nika; 4: On the Double; 5: Windflower); the colors represent the genetic clusters. **C-D**: Relative abundances of the two most abundant taxonomic groups, the collodarians (**C**) and the copepods (**D**). The color gradient and the size of the circles both correspond to the relative abundance.

#### Plankton DNA preservation

The protocol for plankton DNA preservation by heating and desiccation was first tested during two field campaigns along the French Atlantic coast and compared with gold-standard cryo-fixation and preservation. Six citizen crews sampled plankton at different locations (Sup Fig. 4A) using the P2 protocol (Sup Fig. 2, 3). The concentrated plankton samples were equally divided into two subsamples: one was flash-frozen in liquid nitrogen as classically performed on oceanographic vessels for genetics analyses, while the other was heat-dried as implemented in P2 (Sup Fig. 3). On land, samples were preserved in a −80°C freezer (flash-frozen samples) and at room temperature (heat-dried samples) for a couple of months before total DNA extraction and sequencing of 1.3 ±0.07 million V9 SSU rDNA amplicons per sample (Sup. Text 1). Bioinformatic clustering of the plankton communities (defined by types and abundance of rDNA Operational

Taxonomic Units (OTUs, see Sup. Text 1) using different dissimilarity indices indicated that the sub-samples preserved by desiccation and flash-freezing systematically clustered together (Sup. Fig. 4B, C). Plankton communities segregated then first by sampling location, then by their distance to fresh-water input within each bay, irrespective of preservation method. These results proved that the heat drying and subsequent desiccation preservation method do not alter the community composition as measured by DNA metabarcoding and can be applied globally.

#### Empowering pioneer planktonauts to sample the world oceans

With modest seed-funding, we were able to assemble a first set of 20 PlanktoKits, and our primary strategy was to maximize geographic coverage and sampling conditions. We promoted the idea amongst the French sailing community and encountered great enthusiasm to participate. The antique proverb –‘*There are three sorts of people: the living, the dead, and those who sail the sea*’ is profound: mariners are natural engineers, explorers, and planet-lovers. Word of mouth is powerful in their close-knit community and, limited by the low number of sampling kits available, we soon had to start declining requests. The 27 selected pioneer yacht crews, who we call ‘planktonauts’ (Fig. 1A, and https://planktonplanet.org/the-planktonauts/) represented a wide variety of boats and sailing modes, from multi-year expeditions around the world (e.g. Race4Water, Taravana, Folligou), to family cruises (e.g. Manevai, Nika, or Zigomar - see kidsforsea.over-blog.com/), recreation sailing in the same zone over the year (e.g. Suhail), explorers of the poles (e.g. Vagabond or Podorange), or participants in a New Zealand yachting rally across the South Pacific Islands (Fig. 1B, bottom-right insert).

The planktonauts were trained to perform the P2 protocol (Sup. Fig.3), individually or in small groups, and regular internet dialogues were established with them during their voyage to answer their questions and follow their progress. They regularly sent movies and pictures (Fig. 1A, https://planktonplanet.org) of their actions at sea, allowing us to improve our training protocols and outreach. Their feedback on all steps of the protocol at sea (Sup. Fig. 3) have been key to identify the bottlenecks and challenges to overcome for the implementation stage of the scientific program (see below).

#### Geographical coverage and cost of P2-pilot samples

Despite the relatively low number of samples recovered per boat (2 to 18), the widespread routes of many planktonauts (Fig. 1B) yielded plankton samples from 258 sites across the world surface oceans in less than 1 year (Fig. 2B and Sup. Fig.5). This is remarkable compared to the few tens of spatio-temporally relatively restricted stations usually sampled during classical oceanographic cruises, or even to the 147 sites sampled by the schooner *Tara* in 3 years during her first circumglobal expedition (*40*) (Fig. 2B). Furthermore, the total cost of ∼200 US$ per sample (including the price of the kit, training, sampling, shipping, and DNA extractions), which could be significantly reduced with increasing sampling frequency, is one to several orders of magnitude lower than sample cost on oceanographic vessels. The running budget of an open oceanographic research vessel is on the order of 30,000 US$/day. Therefore, a single research vessel traveling at 10 knots and stopping for one hour at each sampling station would have taken 8,6 months - and thus a minimum cost of 7,9 million US$ - to cover 56% of the total P2 sites sampled in slightly more than a year by the planktonauts (Sup. Fig.5). Note that the average cost to equip and run a sailing boat for a transoceanic route is ∼25,000 US$; all together, volunteer planktonauts thus offered ∼90% of the cost of field work in sailing charges. This being said, we acknowledge that planktonauts are logistically constrained and will never be able to gather the sort of comprehensive bathymetric, oceanographic and physico-chemical data that oceanographic vessels routinely collect. P2 is not a substitute, but a valuable complement to the intensive research cruise manned by experts.

### High-quality data for global plankton biodiversity and ecology

18 months after the launch of P2, we had extracted total DNA from 214 plankton samples collected by 27 boats, PCR amplified rDNA metabarcodes from each sample, and generated a total of ±453 million rDNA reads to assess the diversity of eukaryotic plankton (>20 µm) in the explored surface water masses. The methods used for DNA extraction, sequencing, and analyses (Sup. Text 1) were essentially developed in *Tara* Oceans (*34, 52*); the metabarcode used (V9 SSU rDNA) has proven successful to measure the ecological diversity of total eukaryotic plankton (*34*), focus eco-evolutionary analyses on specific groups (*53*–*56*), reconstruct plankton ecological networks (*1*), or revisit plankton macro-ecological patterns (*35, 57*), or biogeochemical processes (*2*).

#### Integration of P2 data into Tara Oceans data. Tara

Oceans (*T*O) data are today a gold standard for ocean plankton ecology, and we first merged the P2 data into the global *T*O metabarcoding dataset (Sup. Text 1) for quality checks and comparison of content. As P2 samples were collected by different sailors, sometimes in harsh conditions, we first checked for the presence of obvious biases, such as the sequencing of bacterial contaminants. After taxonomic assignment of the rDNA reads (Sup. Text 1), we found that the percentage of prokaryotic reads per boat matched the expected value observed in *T*O (given the very large taxonomic spectrum of the eukaryotic PCR primers used that also amplify prokaryotic genes), with just a few outlier samples typically explained by on-board major processing errors (Sup. Fig 6).

We then compared plankton community composition between P2 and *T*O samples. Principal component analysis confirmed the primary influence of organism size on community structuring (*34*), with the pico-nanoplankton (0.8-5 µm) displaying stronger cohesiveness than micro-(20-180 µm) and meso-(180-2000 µm) planktonic communities (Fig. 2A). P2 samples clearly fell within the range of variability of *T*O micro- and meso-plankton samples, reflecting the P2 protocol that uses a 20µm mesh-size net and does not apply any pre-filtration. P2 and *T*O micro/meso-plankton size fractions samples spread together along the second axis of the PCA whose variance (7%) is the result of multiple physico-chemical (dispersal and mixing, variations in temperature, light, nutrients, etc) and biological (species interactions, life cycles, behavior, acclimation/adaptation) processes.

Despite the overall similarity between P2 and *T*O micro/meso-plankton samples (Fig. 2A), the P2 sampling effort did unveil significant novelty in global plankton diversity. All P2 samples produced rDNA OTUs that had not been reported in the *T*O world ocean survey (Fig. 2B), and the number of discovered OTUs clearly increased with sampling effort, both per area (Fig. 2B) and per boat (Fig. 2C). Each planktonaut crew unveiled between 62 and 3,907 unique OTUs, unseen by either *Tara* or any other P2 boats, with a discovery rate partly explained by the eccentricity of the sampled areas relative to *T*O sampling sites. Note for instance the high percentage of boat-specific OTUs (>75%) in the New Zealand area (Fig. 2C).

We further dug into the composition of plankton diversity (rDNA OTUs) unveiled by P2 versus *T*O samples. P2 samples yielded a total of 57,994 OTUs, as compared to 158,716 for *T*O surface ocean samples. Phylogenetic breakdown of the rDNA data (Sup. Fig. 7) shows the overall taxonomic similarity, in both abundance and richness, between data from P2 and the *T*O micro- and meso-planktonic size fractions. While 8,220 OTUs (mostly metazoans and alveolates) were shared exclusively between P2 and the *T*O larger (>20 µm) organismal size fractions, only 1,981 were common with the smaller (<20 µm) *T*O plankton size fractions (Fig. 2D). This emphasizes the consistency of the organisms harvested by P2 and *T*O in comparable plankton size-fractions, and provides organismal data support for our principal component analysis (Fig. 2A). Remarkably, however, more than half of all P2 OTUs (37,163) were actually not seen in any plankton size fractions of the *T*O circumglobal dataset (Fig. 3D). These correspond mostly (∼78%) to Alveolata and Opisthokonta, with an overwhelming majority (98%) of relatively large metazoans (copepods, other arthropods, mollusks, see insert in Fig. 2D). This major difference is most likely explained by the fact that the P2 sampling protocol does not involve an upper-size filtration while the *T*O larger size-fraction was constrained by a sieving at 2mm (micro-plankton: 180-2000 µm), resulting in a more complete survey of larger plankton in P2.

In order to check the value of P2 data at finer-grained taxonomic resolution, we explored the phylogenetic and organismal size-fraction distribution of P2 and *T*O OTUs assigned to a well-known phytoplankton group, the dinoflagellate order Peridiniales (Sup. Fig. 8). Most Peridiniales OTUs observed in *T*O were also found in P2, except a few taxa particularly abundant in the piconano- and nano-size fractions (<20 µm). The relatively large, microplanktonic taxa from the genus *Protoperidinium* were particularly well represented in the P2 dataset, with four OTUs unseen in the *T*O dataset. Note also the presence of many *Blastodinium* and *Brandtodinium* OTUs, which are well-known parasites (*58*) and photosymbionts (*59*) of respectively copepods and radiolarians, and are thus part of the meso- and macro-plankton in their symbiotic stage. Overall, these analyses confirm the quality of the P2 data to assess plankton diversity from the OTU to the community level.

#### Insights into multiscale plankton ecology using P2 data

We finally used P2 metabarcoding data to explore the consistency of macroecological patterns at both global and local scales, across the data collected by the different boats and planktonauts who sampled plankton independently. At the world ocean scale, the P2 data displayed an increase in alpha diversity (Shannon index) from both poles to the tropics (Fig. 3A), reflecting the Latitudinal Diversity Gradient that is well known from terrestrial and marine ecosystems (*60*), and has been previously recorded in several more restricted planktonic groups (e.g. *61*–*63*)) and across plankton kingdoms (*35*). On a more regional scale, the five planktonaut crews from New Zealand collected enough samples to highlight a biogeographical pattern in this area. Clustering of the plankton communities based on their OTU composition into three groups fitted with the three legs of the navigation loop: New Zealand - Fiji; Fiji - New Caledonia, New Caledonia - New Zealand (Fig. 3B). The first area was characterized by high abundances of copepods (Fig. 3D) and important richness of indicator OTUs (*64*) amongst Marine Alveolates (MALV), Acantharia, Dinophyceae and also copepods (Sup. Fig. 9A). By contrast, the last portion of the navigation loop (genetic cluster 3) was characterized by low abundances of copepods and high abundances of collodarians (Fig. 3C), with indicator OTUs belonging to collodarians and spumellarians, diatoms (Bacillariophyta), copepods and choanoflagellates. Group 2 (Fiji-New Caledonia) had a lower diversity of indicator OTUs and was genetically less homogeneous (Sup. Fig. 9B), which may be due to the variability of habitats characterizing this island region. All boats (except *Nika* who is represented by one sample in the analysis) have sampled plankton in at least two different eco-genetic regions, indicating there was no apparent bias linked to the multiplication of samplers.

### Contextual environmental data

Given the importance of the abiotic environment in structuring planktonic ecosystems (*3, 4*), we also included the measure of basic environmental parameters at each P2 sampling site. Besides UTC date/time for each sampling event, the planktonauts were asked to record surface water temperature using the water temperature sensor of their boat and/or mercury thermometer (Fig. 3A), and broad-band spectral water reflectance using the Hydrocolor App (*65*). *In situ* temperature data were compared with those from the NASA’s MODIS-Aqua satellite, and a good match – on average better than 1°C – was observed. We also developed automated procedures to extract remote sensing ocean color data at each sampling site (https://github.com/OceanOptics/getOC), providing bulk information on chlorophyll-a concentration, as well as suspended dissolved and particulate materials in the target water. In the future, this data can not only assist planning of sampling stations at sea in near-real time, but also allow analyses linking ocean color to phytoplankton functional type measured by DNA metabarcoding. On the other hand, the Hydrocolor App has been compared to commercial instrumentation and found to correlate well (*66*); it is hoped it will help relate water reflectance to *in-situ* plankton communities, increasing further the utility of remote sensed ocean color.

## Plankton Planet: perspectives

### Operational PlanktoKit integrating eco-morpho-genetic data

In the pilot stage of ‘Plankton Planet’, we have witnessed the enormous desire of sailors to sample ocean life during their voyages around the globe. We have also generated what is, to our knowledge, the first citizen-based homogenous planetary dataset to explore plankton biodiversity and ecology. Our objective is now to optimize the P2 sampling protocol and tools for large-scale deployment on navigation loops and transects crossing critical ecological and geographical dimensions of the Earth system. Along the way, we have identified critical challenges to overcome.

#### Sampling total plankton at sailing-speed

In the P2 pilot project, the planktonauts were asked to maneuver their boats at a speed of less than 2 knots when towing the 20μm-mesh size plankton net. This requests uncomfortable sailing operations impacting the cruising speed, and it was identified as the primary limiting factor for denser sampling. We have therefore been working on the design of new miniaturized *high-speed nets*, inspired by our successful experience during the *Tara* Pacific expedition (*67*), as well as simple manual pumping systems allowing aspiration of seawater at cruising speed, followed by filtration through a small net system installed on board. The latter device has the advantage of being able to collect pristine water that can also be used to extract DNA/RNA from the smaller plankton size fractions enriched in bacteria, archaea and viruses. Indeed, long-term monitoring of marine plankton systems will ultimately require a sampling protocol covering the 8 orders of plankton organismal size-magnitude, from viruses to animals (*23, 33*), in order to assess both top-down and bottom-up ecological mechanisms shaping the ecosystem.

#### Collecting morphological and behavioral plankton data at sea

The revolution in environmental DNA/RNA sequencing provides the power to comprehensively assess plankton taxonomic and metabolic diversity (*33*). However, meta-omics data convey relatively poor information on the shapes, structures and behaviors of cells and organisms, which may well be the primary drivers for the self-organization of contemporary ecosystems and their emergent functions (*68*). Fundamental phenotypic mechanisms such as symbioses *sensu lato*, selective feeding, or vertical motions and migrations (*69*), and several other unknown complex cellular and organismal behaviors, need to be identified and quantified in the context of the seascape if we ever want to reach a mechanistic pheno-genomic understanding of ecosystem patterning and functioning in the ocean.

Over the last few years, we have thus developed the *PlanktonScope* (see companion paper by Pollina et al., this issue (*51*)), an affordable, miniaturized, modular and evolvable, open source imaging platform for citizen oceanography. For a total cost of less than 400 US$ in parts, the *PlanktonScope* allows both quantitative imaging of plankton communities through a fluidic module before their storage for total DNA/RNA extraction and genetic analyses in the lab, and high-quality imaging/filming of individual cells or organisms under different types of illumination. The scarcity of live images and movies of plankton at sea constitutes arguably the major knowledge-gap in oceanography as today’s oceanographers spend most of their time on land behind their computer screens or analyzing dead plankton in the lab. The *PlanktonScope* will not only bring planetary imaging of plankton life through its global seatizen deployment, but also, we hope, trigger an emotional shock in each planktonaut discovering the vibrant beauty of the marine microbiome, changing their view of the ocean forever and further driving their will to explore and preserve the invisible world thriving under their boat.

### Toward a perennial, self-sustainable standardized citizen monitoring of global ocean life

The cost of the upcoming P2 PlanktoKit 1.0 (plankton collecting system, PlanktonScope, DNA-kit, software App to visualize, integrate, and store eco/morpho/genetic information) is estimated at 2,000-3,000 US$ per boat, allowing its deployment on hundreds of seatizen boats. The deployment of the first 100 kits will start in 2021 in the Atlantic and Pacific oceans, and is designed to address topical questions in ocean ecology and to test current models and theories that predict patterns of ocean biodiversity and functionality in relation to the physics of marine water and climate.

Based on feedback between these unique data and modelling efforts, we will then fine tune the optimal density of sailboats (or fish boats or cargo ships) and sampling frequency for longer-term deployment of the P2 protocol on key navigation loops and routes that cover the world oceans. Our ultimate goal is to implement, over the next decade, a self-sustainable and standardized global survey of the diversity and evolution of surface ocean plankton ecosystems. We believe that this endeavor can only happen through collaboration at the interface between the worlds of sailors, makers, and researchers, and it can fundamentally change the way oceanography is performed, while enhancing the collective knowledge and consciousness of the open ocean invisible life at the core of global ecology (Box 2).

#### BOX 2.

**Perspective outputs of Plankton Planet**

##### Oceanography 3.0 ecosystem

- An international fleet of planktonauts to act as sentinels and the collective consciousness of the biological health of our oceans.
- An evolvable toolkit of affordable, scientifically relevant instruments for seatizen-based assessment of aquatic (marine and freshwater) biodiversity and ecosystems.
- An ever-growing cryo-bank of global ocean DNA samples, archiving the memory of our changing oceans for future generations and technologies.
- A continuous flow of standardized ocean imaging and genetic data at unprecedented spatial, temporal, and taxonomic scales, freely available for fundamental and applied science, policy-makers, and education.

##### Blue-sky science

- Exceptional long-term monitoring of the distribution and evolution of total plankton biodiversity in our fast-changing ocean; including analysis of the global impacts of warming, acidification, and de-oxygenation.
- Novel understanding of abundance, structures, functions, and behaviors of plankton life via the billions of images and movies of planktonic organisms generated using PlanktonScopes.
- Incorporation of high-resolution global-scale comprehensive biological data into efforts to model the dynamics of ocean ecosystems and ocean-climate interactions, from genes to biogeochemical cycles.

##### Applied science & policies

- Detection of invasive, toxic, or economically relevant species at a planetary scale.
- Linking of data on plankton communities (including fish gametes and larvae) to fish-catch data for robust prediction of fish stocks.
- Assessment of ocean biogeographic zones (“seascapes”) for optimal design of marine protected areas in the high seas (e.g. ecosystems with high capacity for carbon pumping).
- Evaluation of the health of oceanic regions based on species content, species richness, and trends in time.

##### Education through ‘Plankton & Arts’

- Collective awareness of oceanic microbiomes and their planetary impact, both directly (through the PlanktonScope) and indirectly via the shared images, movies, and 3D prints of plankton. Educational tools will notably be distributed by planktonauts in remote countries and islands whose communities interact with and depend on the marine ecosystem.

### Citizen ecoscopes for collective field assessment of the biosphere

At the core of our long-term vision, the principles applied to development of the PlanktonScope (*51*) will be extended to co-construct a field-EcoScope for citizen exploration of aquatic eco-systems, and the concept could be easily extended for the exploration of other planetary biomes. Like the PlanktonScope, the field-EcoScope is a miniaturized platform that can host *modules* and sensors to measure the critical biological, but also (bio)chemical, and physical parameters of any eco-system. Each module (including the PlanktonScope) will be co-developed by small international and interdisciplinary teams of (*i*) researchers who will define the scientific specifications to address fundamental question in global ecology, (*ii*) mariners and citizen samplers who will assess practical user-constraints linked to specific field conditions (e.g. *on-board* sailing boats or cargo-ships), and (*iii*) makers and freelance engineers who will design and construct the frugal modules to measure given parameters. The Plankton Planet ecosystem will manage communication and provide the overall coordination to ensure that all modules can be fully integrated into the miniaturized field-laboratory and generate good-quality standard data that are immediately shared in public databases.

We believe that it is a matter of urgency to quantify and understand the structure, dynamics, and evolution of *eco-systems*. Over the last century, societies have spent phenomenal amounts of money to discover the properties of the infinitely small (atoms and cells) and the infinitely large (weather, planets and stars, and the universe). But it is in between these extremes that we find arguably the most complex object in the known universe: eco-systems, i.e. self-organized and evolving life networks that interact with physico-chemical processes at micro-to planetary scales. Ecosystems have shaped the Earth atmosphere and biogeochemical cycles, they currently buffer climate change while providing sources of food and medicine, they will determine the future habitability of our planet and the fate of humanity. Due to their extreme complexity, integrating biology, chemistry, and physics, ecosystems have long escaped holistic quantitative assessment. Today the tools and methods exist, in particular automated sequencing and imaging, as well as artificial intelligence and massive computing to collect and integrate the layers of big data needed to understand eco-systems. The major challenge is to develop the new generation of affordable tools that can be deployed in a systematic manner across the spatio-temporal dimension of the Earth system. Plankton Planet proposes a coherent and frugal approach to overcome this challenge, at the interface of science and society, using one of the simplest biomes – marine waters-as a case study. Clearly being at the onset of planetary biology, we hope that P2 will contribute to reaching a profound understanding of our habitat in the decades to come, and to learning how to live in synergy with the biosphere.

## Acknowledgments

We first deeply acknowledge the generosity and curiosity of the pioneer planktonauts who have collected plankton samples from the world oceans and made the P2 proof-of-concepts possible: E. Abadie on board *Manevai*, I. Autissier on board *Ada 2*, R. Barnaud on board *Kauana*, C. Beaumont on board *Folligou*, E. Brossier on *Vagabond*, K. & K. Brownie on board *Astral Express*, V. Chirié on board *Dreamweaver*, I. Corrias on board *Attila 3*, Q. d’Avout on board *Vitamin Sea*, B. de Ravignan on board *La Cardinale*, M. Denis on board *Isbjørn II*, E. Di Iorio on board *Suhail*, B. Dumontet on board *Ainez*, E. Dupuis on board *Vaihere*, N. Fabbri on board *Zigomar*, Y. Gladu on board *Tethys*, J. Goulias on board *Gwalarn*, M. Hardy & A. Schnarwiler on board *Taravana*, P. Laparre on board *Amasia*, A. Marchandise on board *Karukera*, L. Marie on board *Vagabond*, J. & L. Martin on board *Windflower*, C., L., G., O. & R. McIntyre on board *Nika*, M. & C. Stephens on board *The Double*, B. Monégier on board *Podorange*, D. & S. Patterson on board *Celine*, Lorena Piana on board *Giulia*, A. Schmid on board *Julo*, M. Simeoni on board *Race for Water*, Students on Ice on board the MV Ocean Endeavor, Tara’s crew on board *Tara*, S. Valcke on board *Manamo*, & JM. Viant on board *Dame Jane*. Thanks also to all members and volunteers of the P2 team: Romain Bazile, Carole Beaumont, Emmanuel Boss, Guillaume Bourdin, Barry Cael, Marie Caillaud, Roberto Casati, Sébastien Colin, Johan Decelle, Christelle Demonte, Colomban de Vargas, Mick Follows, Gaby Gorsky, Damien Guiffant, Jeremy Guyard, Nils Haëntjens, Nicolas Henry, Rainer Kiko, Aurélie Labarre, Adam G. Larson, Noan Le Bescot, David Le Guen, Fabien Lombard, Frédéric Mahé, Gilles Mirambeau, Clémentine Moulin, Anna Oddone, Xavier Pochon, Thibaut Pollina, Manu Prakash, Ian Probert, Sarah Romac, Romain Troublé, Flora Vincent, Marie Walde. The Plankton Planet Pilot Project was made possible by seed-funding from the Richard Lounsbery Foundation. Additional funding came from the French Government “Investissements d’Avenir” program OCEANOMICS (ANR-11-BTBR-0008), the ‘*Plankton Arts*’ grant from the Fondation d’Entreprise Total, the New Zealand Royal Society Dumont-D’Urville (DDU-CAW1501) and Strategic Seeding (16-CAW-008-CSG) Funds, the Cawthron Institute, as well as the Okeanos, *Tara* Ocean, and Schmidt Futures Foundations. We are grateful to Pierre Mollo, Francoise Gaill, Eric Karsenti, Paul Falkowski, Étienne Bourgois, Catherine Chabaud, Roland Jourdain, Isabelle Autissier, Raphaela le Gouvello, Daniel Richter, Chris Cornelisen and Gilles Boeuf, for their continuous advice and support. This article is contribution number 1 of Plankton Planet.

## P^2^ on the web

- Web site: http://planktonplanet.org

- YouTube channel: http://youtube.planktonplanet.org

- Vimeo channel: http://vimeo.planktonplanet.org

- Twitter account: https://twitter.com/PlanktonPlanet (@PlanktonPlanet)

---

**Supplementary Figure 1.**
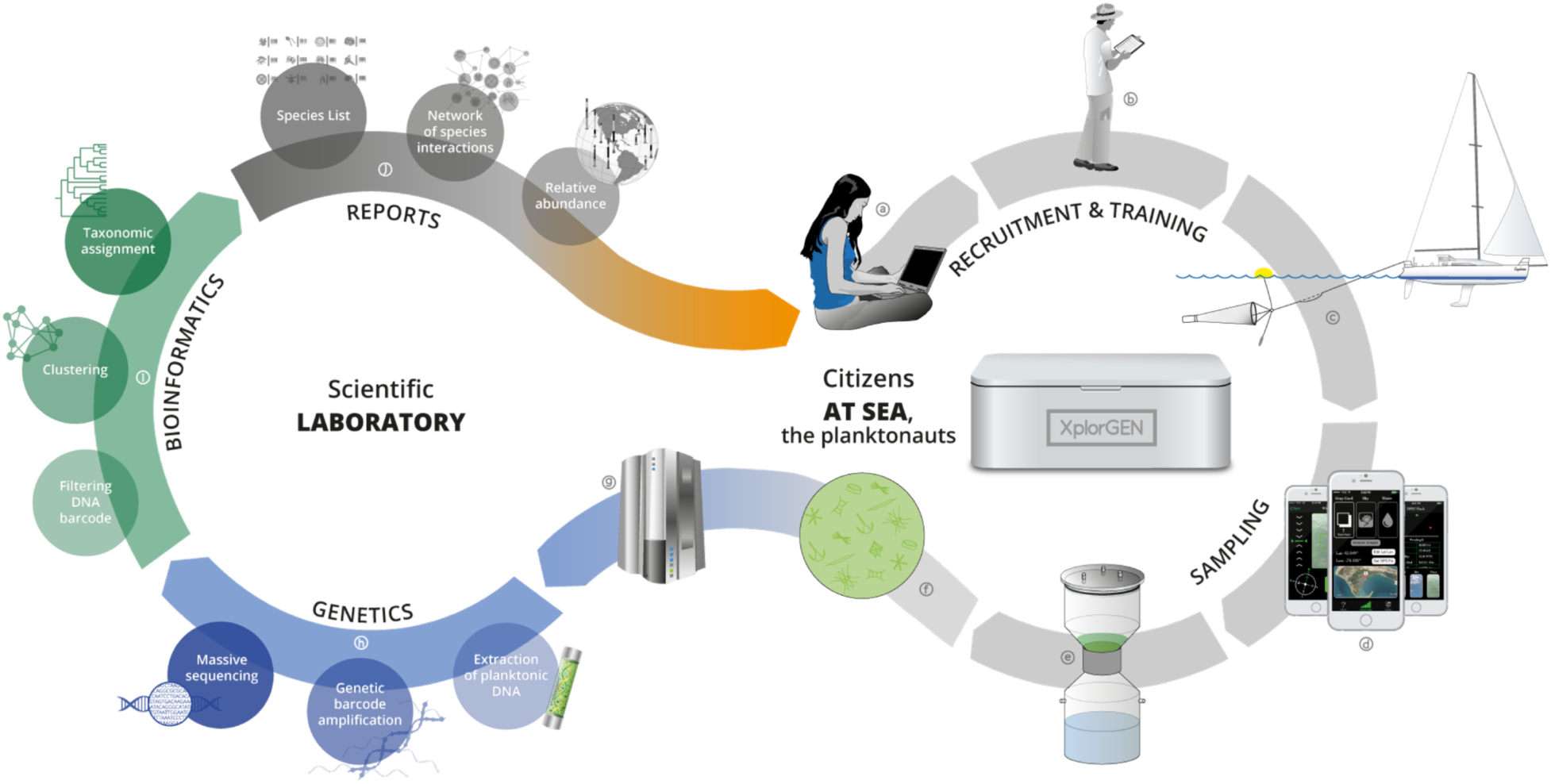
**Original Plankton Planet strategy,** showing the successive steps of both the seatizen (right loop, a to f) and researchers (left loop, g to j) operations, respectively at sea and in the lab.

**Supplementary Figure 2.**
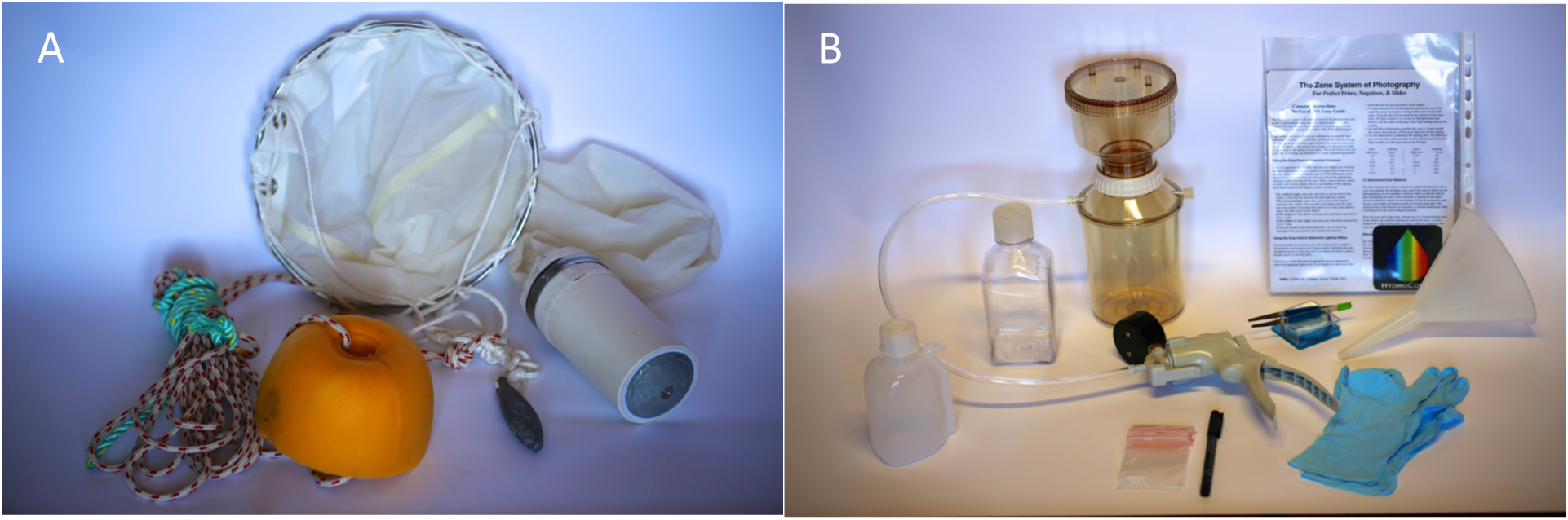
The original Plankton Planet sampling kit. A. The P2 plankton net (25cm diameter, 20μm mesh size) with a 2kg weight and a small float to maintain it at a maximum depth of 3m. B. The manual vacuum pumping system used to transfer plankton from the cod-end of the net onto a 10μm filter membrane. The total cost for one kit approximates 700 US$ but could be largely reduced for mass production via tinkering of home-made parts in FabLabs.

**Supplementary Figure 3.**
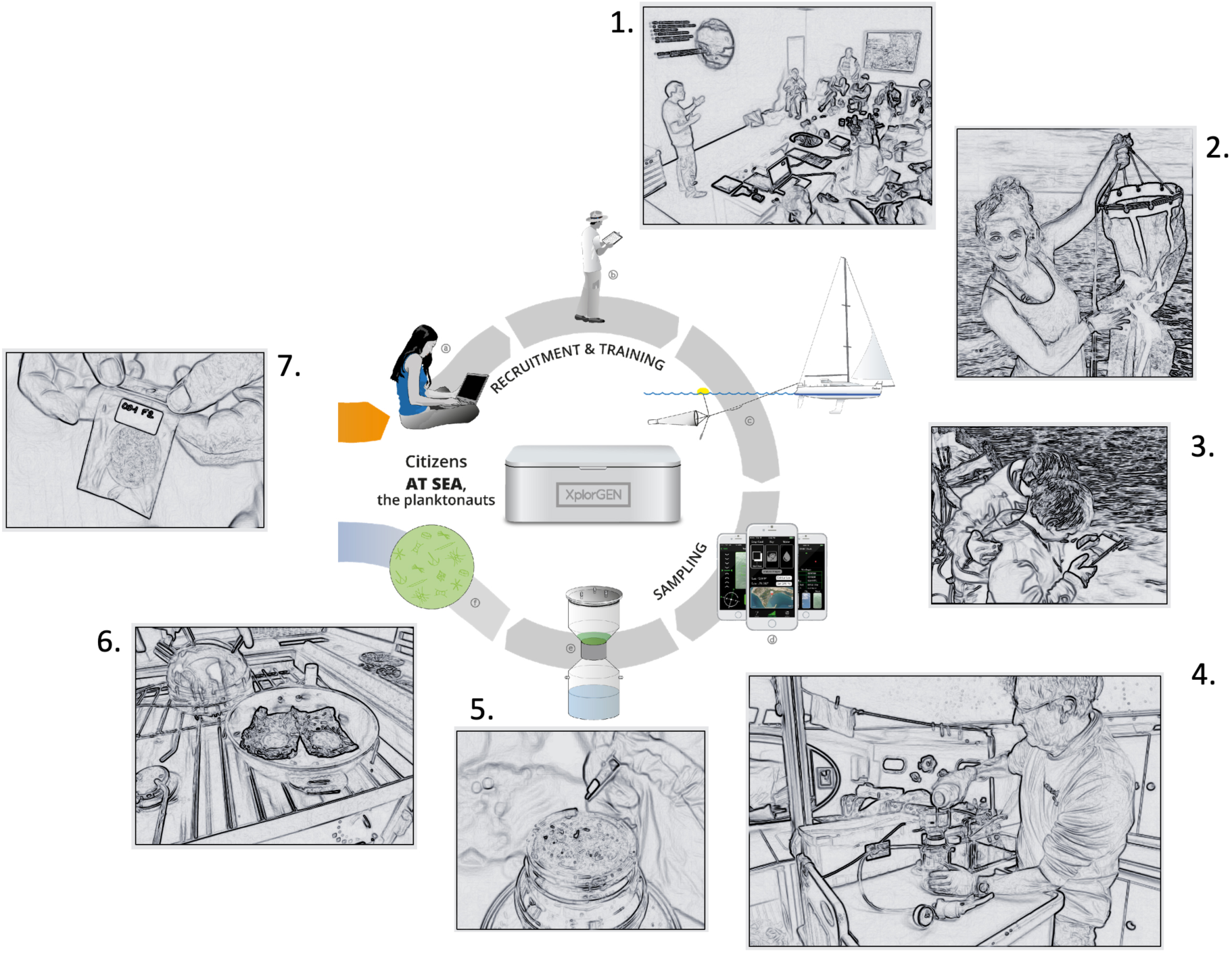
Pictures illustrating various steps of the sampling protocol for planktonauts at sea. 1. Training of a group of planktonauts, here in Auckland with Dr. Pochon. 2. After 15-20min of towing at a maximum speed of 2 knots, the plankton net is recovered on board, here onboard *Tethys*. 3. Recording of contextual parameters, here ocean color using the *Hydrocolor App* on board *Zigomar*. Note that 3 families participated to the pilot project, showing that even kids can realize parts of the protocol (see for instance: https://vimeo.com/219660346); 4. Pouring of the concentrated plankton from the net cod-end into the manual vacuum-pumping, onboard *Taravana*. 5. Manual recovery of the 10μm filter membrane full of plankton biomass. 6. Gentle drying of the filter membrane in a pan on the boat (*Taravana*) gas cooker. 7. Storage and labeling of dried plankton samples into zip lock plastic bags before shipping to the laboratory by regular mail.

**Supplementary Figure 4:**
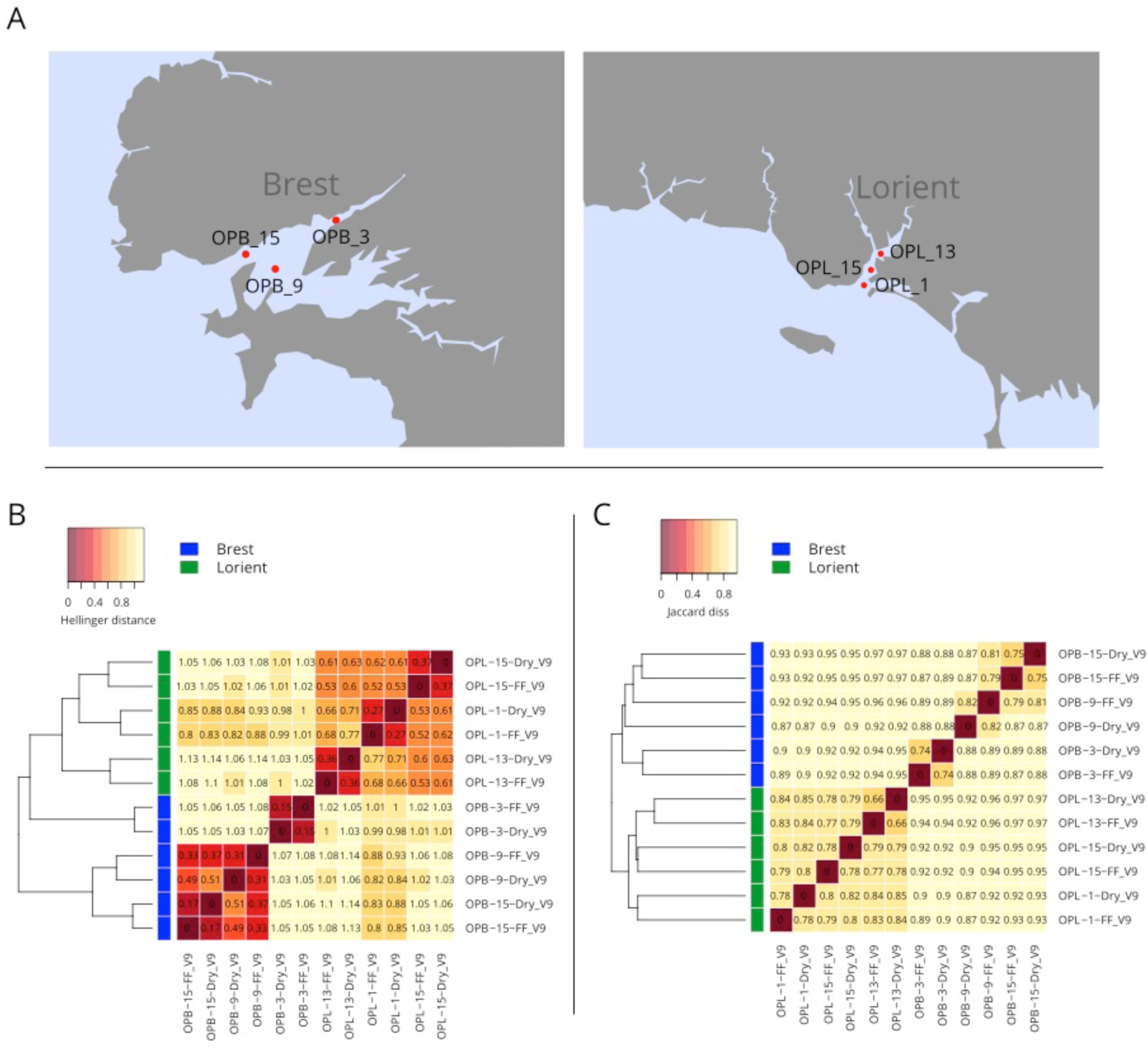
Testing the effect of dried (P2) versus flash-frozen plankton samples on DNA-based community composition. A. Geographic location of plankton samples collected by volunteer citizen sailors along the French coast during the ‘Objectif Plankton’ initiative (https://riem-asso.com/objectif-plancton/). Bay of Brest, June 21^st^ 2016, on the left, and Bay of Lorient, June 14^th^ 2016, on the right; OPB and OPL indicate samples from Brest and Lorient bays, respectively. Each sample was preserved by both desiccation (Dry) and Flash Freezing into Liquid Nitrogen (FF), before deep Illumina sequencing of V9 SSU rDNA amplicons. B. Heatmap of the Hellinger distance (Hellinger standardization followed by Euclidian distance) between samples and sub-samples. C. Same as B, using Jaccard distances (based on OTUs presence/absence). The relatively high values of the Jaccard distances between preservation methods from the same samples comes from the fact that low abundance OTUs create a noisy background that inflates dissimilarities. The Hellinger distances are less sensitive to this issue, and the dissimilarities values range from 0.15 to 0.37 between preservation methods, while they are close to 1 between samples. The residual dissimilarities observed between subsamples are due to Illumina sequencing errors and are expected between technical replicates.

**Supplementary Figure 5:**
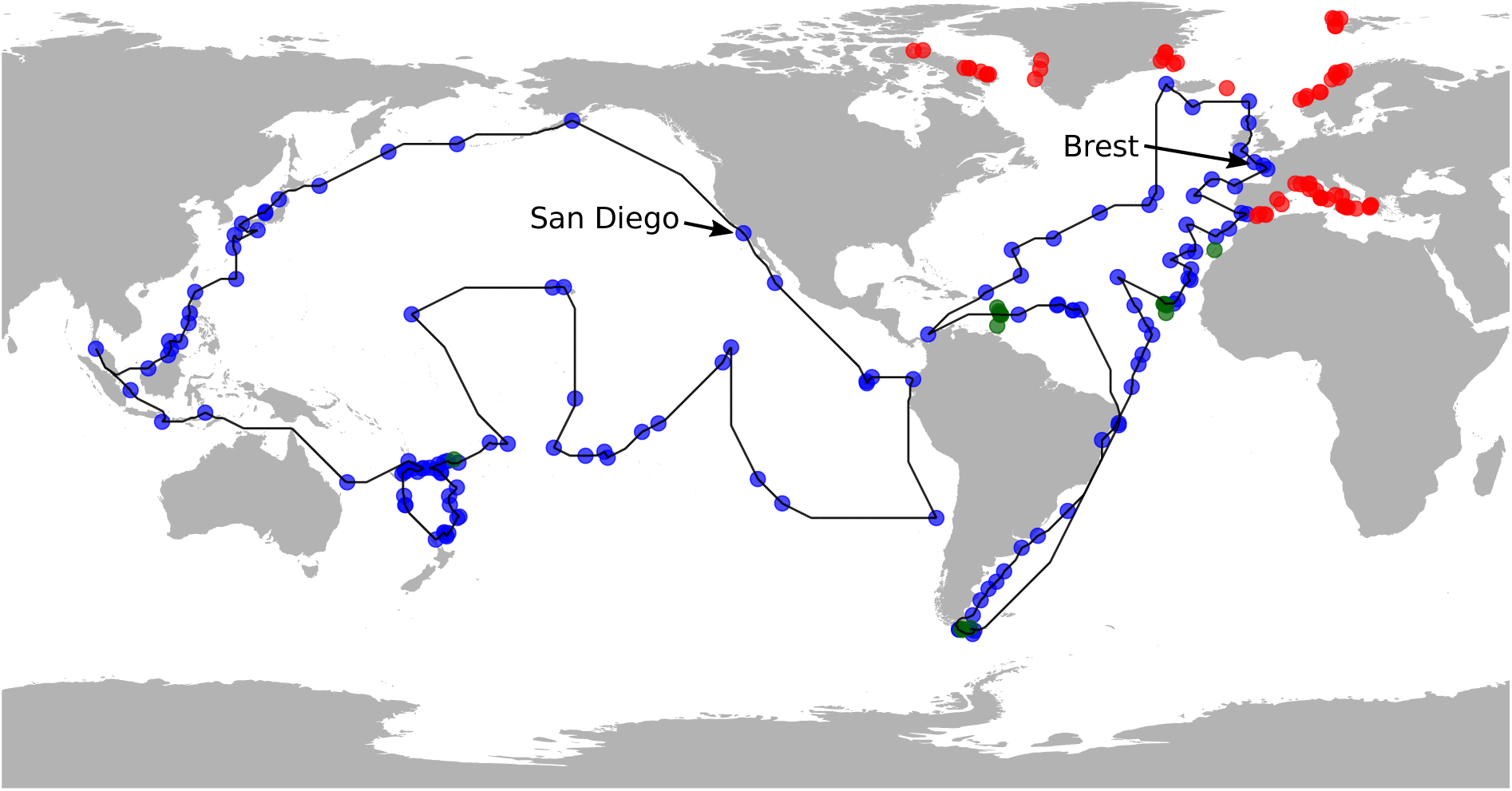
Geographical coverage and putative cost of P2-pilot samples. Location of 233 (out of 258) ocean sites sampled by the pioneer planktonauts in 14 months during the Plankton Planet pilot project. The blue dots are the sampling sites taken into account to compute the shortest possible route by a putative single oceanographic vessel in the Indo-Pacific and Atlantic basins, respectively (‘traveling salesman problem’, see Sup. Text 1). The green dots are sampling sites considered as land sample (depth >0 m) because of their proximity to islands and the lack of resolution of the bathymetric database used to compute the shortest path without crossing the land between samples (see Sup. Text 1). The red dots in the Mediterranean sea and Arctic ocean, correspond to sampling sites that were not integrated into the ‘traveling salesman problem’. The Atlantic loop, starting and ending in Brest, pass through 62 sites and is 44,972 km long; the Pacific loop, starting and ending in San Diego, pass through 82 sites and is 68,985 km long. Using a conventional oceanographic vessel (30,000 $US a day) travelling at 10 knots and stopping at each station for 1 hour, sampling these two loops would cost circa 7,9 million $US in total.

**Supplementary Figure 6:**
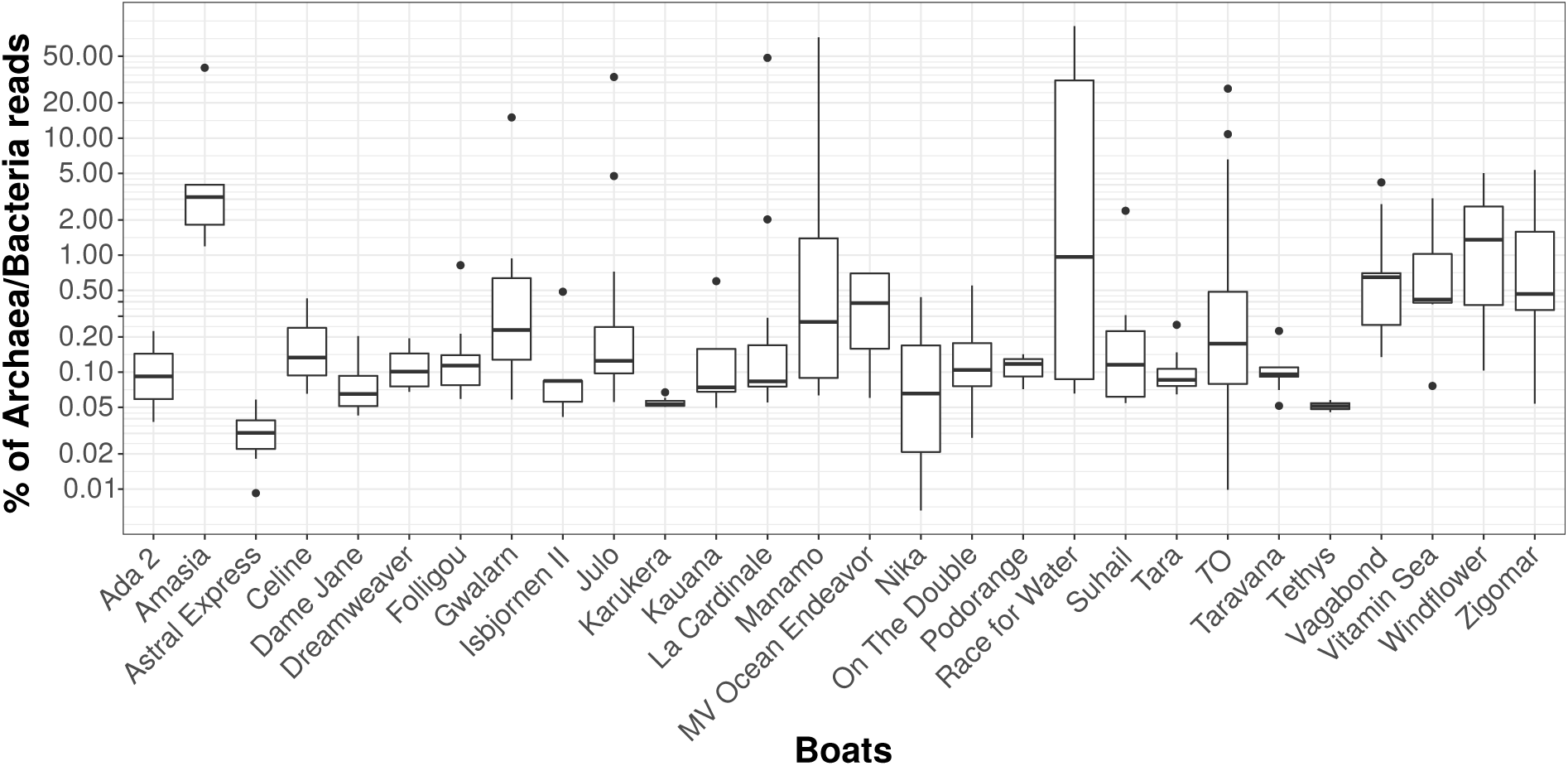
Box plot representation of the ***percentage of prokaryotic (Archaea and Bacteria) rDNA reads P2 samples*** for each boat, and for *Tara* Oceans samples (micro- and meso-planktonic size fractions). 85% and 89% of P^2^ and *T*O samples, respectively, contain less than 1% of prokaryotic reads, the difference being explained by a few outliers. Note the higher value for a few samples from *Amasia* and *Race for Water*, which correspond to samples that were kept on board at room temperature for several days before processing.

**Supplementary Figure 7:**
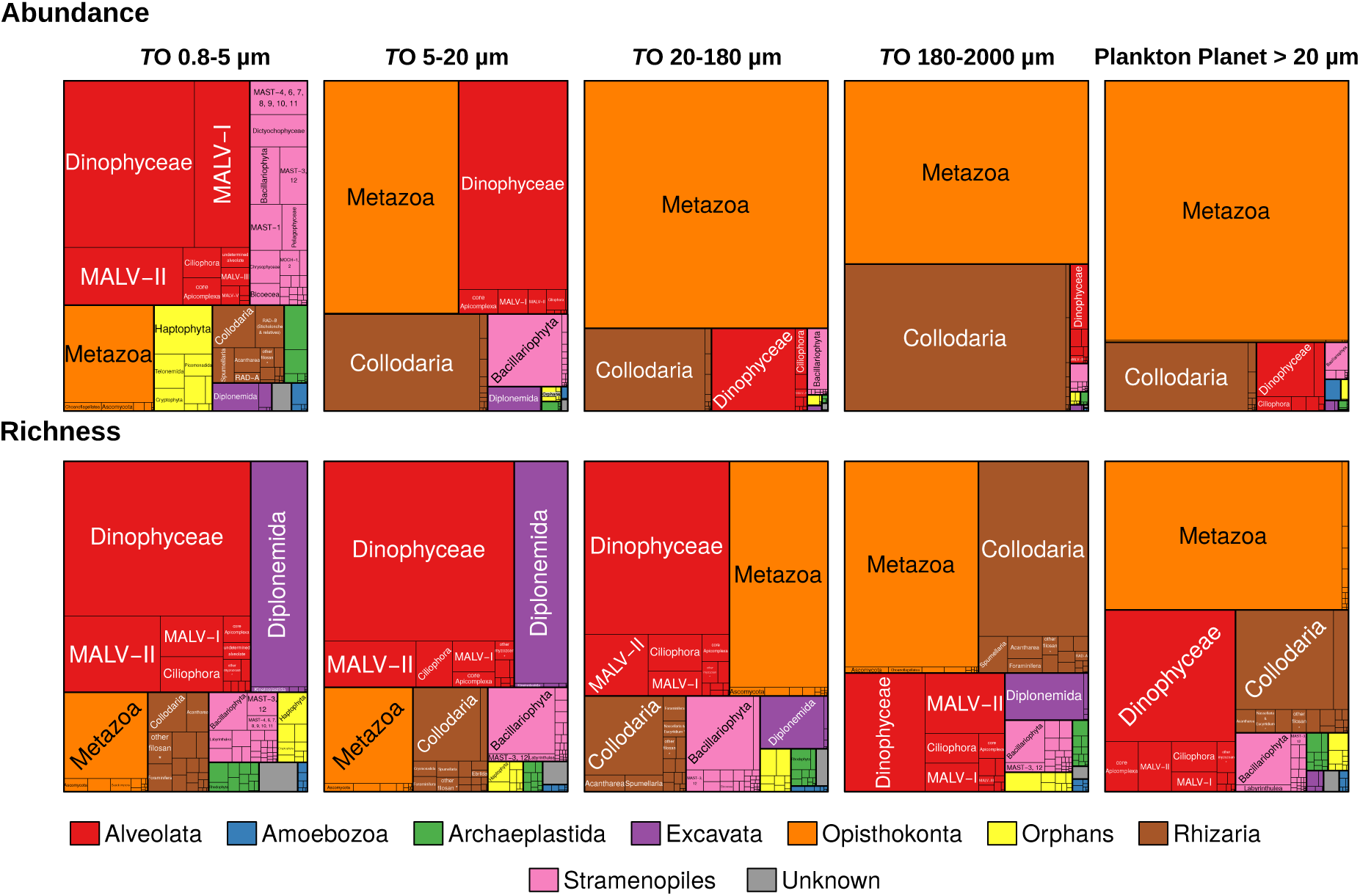
Phylogenetic breakdown of the Tara Oceans and Plankton Planet global ocean metabarcoding datasets at the eukaryotic supergroup and ‘taxogroup’ (sensu^1^) levels. All V9 rDNA reads and OTUs with genetic similarity to a eukaryotic reference sequence ≥80% were retained and taxonomically assigned. The tree-maps display the relative abundance (upper part) and richness (lower part) of the different taxonomic groups in *T*O plankton size fractions and P2. The category ‘*Orphans’* contains the known but phylogenetically uncertain deep-branching lineages (i.e. Haptophyta, Telonemida, Picomonadida, Katablepharidida, Cryptophyta, Centrohelida and Apusozoan); ‘*Unknown’* corresponds to OTUs assigned to two different supergroups.

**Supplementary Figure 8:**
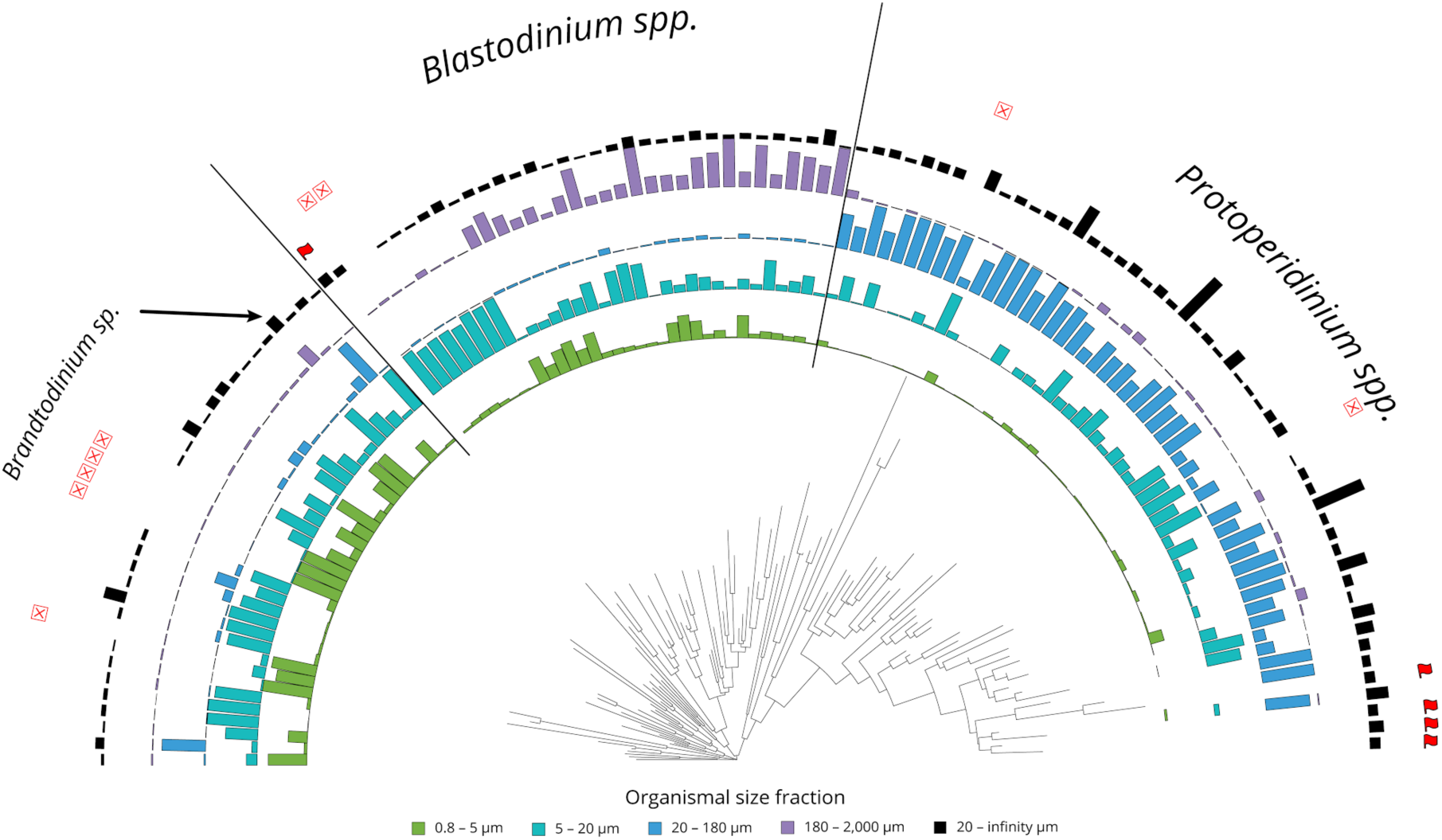
Phylogenetic and plankton size fraction distribution of the most abundant OTUs assigned to Peridiniales dinoflagellates in both the P2 and TO metabarcoding datasets. The 100 most abundant Peridiniales OTUs in TO and the 50 most abundant in P2 were selected, together representing 119 unique OTUs. Colored bars indicate the proportion of mean relative abundance of each OTU amongst the four *T*O plankton size fractions. Black bars indicate the mean abundance (log transformed and scaled to 1) of each OTU amongst all P2 samples. Flag and red crosses symbols indicate OTUs observed respectively only in P2 and only in TO.

**Supplementary Figure 9:**
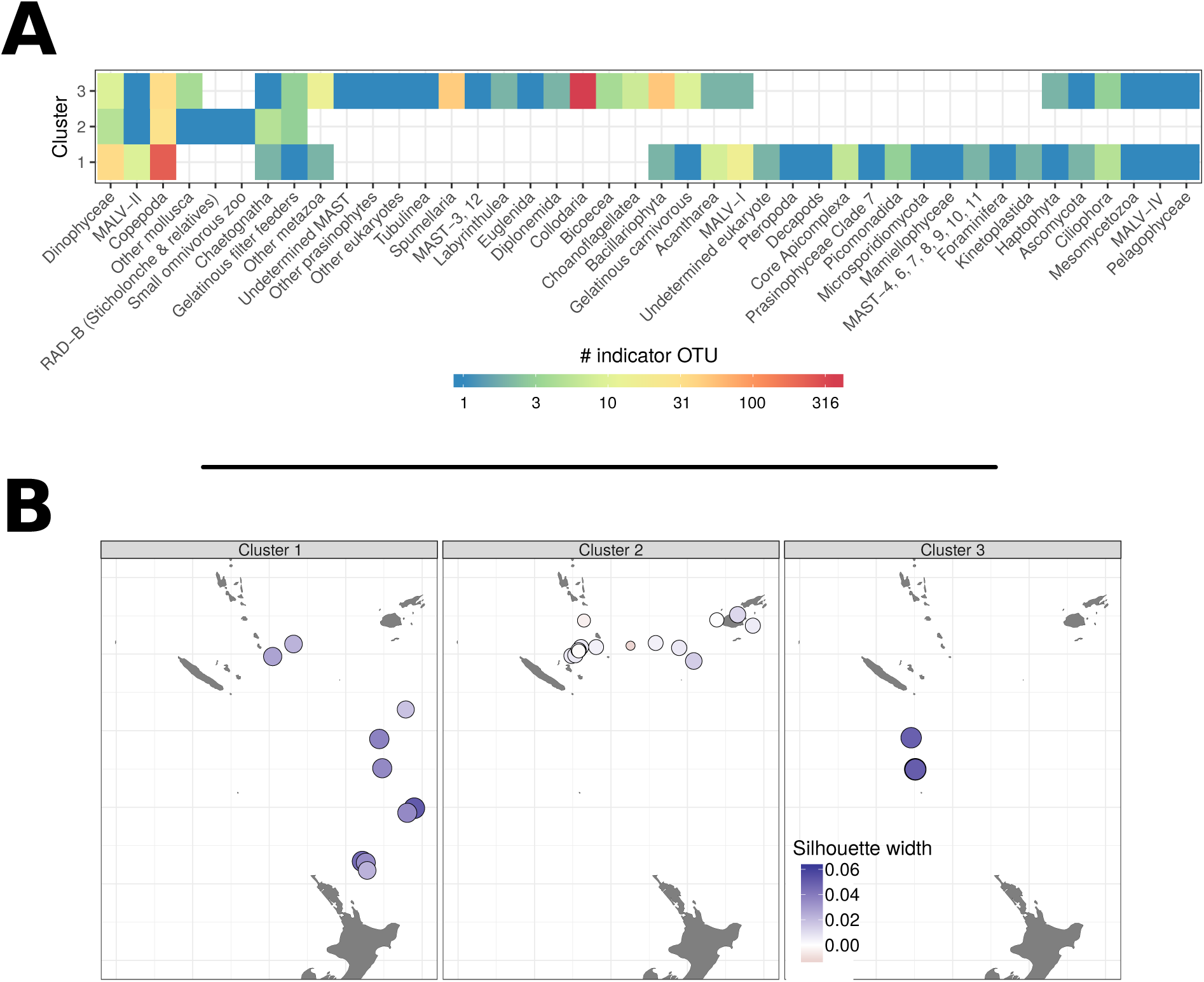
Details about the three plankton community clusters generated for the South-West Pacific loop samples (Fig. 4B). **A**: Geographic distribution of indicator OTUs among taxonomic groups for each cluster. For each OTU, the indicator values, i.e. the product of the relative frequency and relative average abundance in clusters, are computed from Hellinger transformed abundances for each of the three clusters using the indval() function (labdsv R package, ^49^). OTUs are considered as indicator of a specific cluster when p-value are below 0.01. **B**: Consistency of the clusters using the silhouette width (measure of how similar a sample is to its own cluster compared to other clusters). The silhouette width has been computed using the silhouette() function (cluster R package^43^).

## Supplementary Text 1. Material & Methods

### P2 protocols: plankton DNA sampling, storing, shipping, extracting, and sequencing

After 15-20 min of net towing at a speed of ∼2 knots (Sup. Fig. 2), 500mL of concentrated seawater were split and manually filtered onto two replicate 10µm polycarbonate filter-membranes (diameter 47mm), dried at 70°C for 5 min in a pan (on the boat gas-cooker), and then stored at room temperature in a hermetic Ziploc bag with granular desiccants (Sup. Fig. 2, 3). Upon arrival in a port, samples were shipped to the Roscoff Marine Laboratory (Brittany, France) in a simple envelope via regular mail. In the laboratory, the samples were stored at −80°C and information related to samples was archived in a database together with contextual data.

DNA extraction was performed using a protocol modified from the *NucleoSpin Plant Midi* kit (Macherey-Nagel). One replicate filter of each sample was cut in small pieces and incubated for 2h at 56°C with 3.6mL of the lysis buffer PL1 and 250µL of proteinase K. The lysate was transferred to a large capacity *NucleoSpin* Filter (*DNA Midi kit*) and centrifuged for 10 min at 1,500 g. The eluate was transferred to a new tube and 1 volume of PC buffer was added. The mixture was loaded into the appropriate spin column and washed 3 times with the DNA wash solution. Total DNA was finally eluted twice with 150 µl of DNA elution buffer, and stored in sterile microtubes at −20°C. The amount of recovered DNA was quantified by dsDNA-specific fluorimetry using a Qubit 2.0 Fluorometer with Qubit dsDNA Broad Range and High Sensitivity Assays (ThermoFisher Scientific, Waltham, MA). The DNA quality was double-checked in a subset of samples by running 1 µl on 1,2% agarose gel for 45 min at 120V.

To address general questions of eukaryotic biodiversity over extensive taxonomic and ecological scales, the hyper-variable loop V9 of the Small Sub-Unit (SSU) ribosomal (r) RNA gene was targeted for the generation of amplicons by Polymerase Chain Reaction (PCR). This barcode presents a combination of advantages: (*i*) it is universally conserved in length and simple in secondary structure, thus allowing relatively unbiased PCR amplification across eukaryotic lineages followed by *Illumina*sequencing, (*ii*) it includes both stable and highly-variable nucleotide positions over evolutionary time frames, allowing discrimination of taxa over a significant phylogenetic depth, (*iii*) it is extensively represented in public reference databases across the eukaryotic tree of life, allowing taxonomic assignment amongst all known lineages. Amplifications of the V9 from SSU rDNA were conducted with Phusion® High-Fidelity DNA Polymerase (Finnzymes) using the PCR primers 1389f 5’-TTGTACACACCGCCC −3’ and 1510r 5’-CCTTCYGCAGGTTCACCTAC −3’. The PCR mixture (25µL final volume) contained 5ng of template with 0.35µM final concentrations of each primer, 3% of DMSO and 2X of GC buffer Phusion Master Mix (Finnzymes). Amplifications were conducted following the PCR program: initial denaturation step at 98°C for 30 sec, followed by 25 cycles of 10sec at 98°C, 30sec at 57°C, 30sec at 72°C, and a final elongation step at 72°C for 10 min. Each sample was amplified in triplicate to get enough amounts of amplicons. Products of the reactions were run on a 1.5% agarose gel to check for successful amplification products of the expected length. Amplicons were then pooled and purified using the NucleoSpin® PCR Clean Up kit (Macherey-Nagel, Hoerdt, France), and finally quantified with the Quant-iT™ PicoGreen ® dsDNA kit (Invitrogen). Purified amplicons were sent to Genoscope for HiSeq sequencing.

### Sampling coverage and putative cost

In order to compare ocean sampling cost between P2 sailing boats and classical oceanographic vessel, we measured the shortest possible route between P2 sampling sites (traveling salesman problem) by a putative oceanographic vessel. For convenience, 144 P2 sampling sites (56% of total sites) from the Pacific (83), Indian (1) and Atlantic (60) Oceans with a depth >0m according to the ETOPO1 NOAA database (*70*) were selected. We then considered two ports as starting points for two navigation loops: Brest for the Atlantic Ocean samples plus 2 Pacific Ocean samples located at the extreme South of South America (62 samples in total); and San Diego for the other Pacific and Indian oceans samples (82 samples in total). For each group of stations, the shortest possible route that visits each station and return to the starting port was defined. To avoid crossing lands, the least cost distances between locations were computed by constraining paths to depths below 0 m. The distance matrix was computed using the lc.dist() function from the marmap R library (*71*). From this matrix, the traveling salesman problem was solved for each basin with the arbitrary insertion algorithm followed by a two edge exchange improvement procedure using the solve_TSP() function with default options (TSP R library (*72*)).

### Bioinformatic data processing

In order to compare P2 DNA metabarcoding data to the primary *Tara* Oceans (*T*O) eukaryotic metabarcoding dataset (*34*), we first merged raw reads from both datasets and applied the following bioinformatics steps. Paired Illumina™ MiSeq reads from the 214 P2 samples and the 883 samples from *Tara* Oceans (2009-2012) were assembled with vsearch v2.7.1 (*73*) using the command fastq_mergepairs and the option fastq_allowmergestagger. Demultiplexing and primer clipping were performed with cutadapt v1.9 (*74*) enforcing a full-length match for sample tags and allowing a 2/3-length partial match for forward and reverse primers. Only reads containing both primers were retained. For each trimmed read, the expected error was estimated with vsearch’s command fastq_filter and the option *eeout*. Each sample was then dereplicated, i.e. strictly identical reads were merged, using vsearch’s command derep_fulllength, and converted to FASTA format. To prepare for clustering, samples were pooled and submitted to another round of dereplication with vsearch. Files containing expected error estimations were also dereplicated to retain only the lowest expected error for each unique sequence. Clustering was performed with Swarm v2.2.2 (*75*), using a local threshold of one difference and the *fastidious* option. The representative sequences of each molecular operational taxonomic unit (OTU) were then searched for chimeras with the vsearch’s command uchime_denovo(Edgar et al., 2011). In parallel, the OUT representative sequences were taxonomically assigned with the stampa pipeline (https://github.com/frederic-mahe/stampa/) and a custom version of the Protist Ribosomal Reference database PR2 (*76*). This custom reference database is an update of the V9_PR2 reference database used for the taxonomic assignation of the *Tara* Oceans metabarcodes (*34*).

Clustering results, expected error values, taxonomic assignments and chimera detection results were used to build a raw OTU table. Up to that point, reads that could not be merged, reads without tags or primers, reads shorter than 32 nucleotides and reads with uncalled bases (“N”) were eliminated. To create the “cleaned” OTU table, additional filters were applied to retain only: non-chimeric OTUs, OTUs with an expected error per nucleotide below 0.0002, and OTUs containing more than 3 reads or seen in 2 samples. The final OTU table contained integrated the 214 P2 samples together with the 386 *Tara* Oceans samples collected in surface waters for 4 organismal size fractions (0.8-5, 5-20, 20-180, 180-2000 µm). This table was subsampled (rarefied) at the minimum number of reads observed for a sample (313,539 reads) for comparison of OTU richness between samples. Based on the rarefied OTU table, an OTU is considered as unique to P2 if it is present in at least one P2 sample and not present in any of the 386 *Tara* Oceans surface samples. The plankton samples from ‘*Objectif Plancton*’ used to compare the preservation methods were processed independently using the same bioinformatic pipeline. Raw data are available under the bioproject XXXXX (https://www.ncbi.nlm.nih.gov/bioprojectXXXXX).

